# Assembly of 23 Plastid Genomes Provides the Chloroplast View on *Miscanthus* Origins

**DOI:** 10.1101/2020.06.28.175893

**Authors:** Dyfed Lloyd Evans

## Abstract

Despite its economic importance as a new biofuel resource, little work has been done on the large-scale phylogenetics of *Miscanthus*. Twenty-three complete *Miscanthus* chloroplasts were assembled and annotated. A phylogeny was performed with these assemblies, which shows the relationships between the main *Miscanthus* species and sub-species. The phylogeny demonstrates that there is no meaningful distinction between *Miscanthus floridulus* and *Miscanthus transmorrisonensis* as the accessions are not distinct. However, at the crown of the tree there is a clear distinction between *M. sinensis malepartus* and *M. sinensis condensatus* subspecies. The phylogeny reaveals that the female parent of *Miscanthus xgiganteus* is *Miscanthus lutarioriparius* rather than *Miscanthus sacchariflorus*. The phylogeny also shows a novel hybrid between *Miscanthus oligostachyus* and *Miscanthus sinensis*, a grouping to which *Miscanthus sinensis* var Purpurascens belongs. This hybrid form is named *Miscanthus* ×*oligostachyus*.

## Introduction

Since the 1970s, *Miscanthus* has come to the attention of plant breeders as a potential source of bioenergy and fibre (Jones and Walsh 2001). The genus Miscanthus comprises perennial rhizomatous grasses with a predominantly tropical distribution (Clayton and Renvoize 1986). However, members of the genus occur from Mediterranean Europe and North Africa through tropical Asia to New Guinea, Japan and temperate China (Clifton-Brown et al. 2008) and a centre of diversity in Temperate Asia.

Despite the interest in some hybrids and species of *Miscanthus* as energy crops, these are predominantly clonally propagated and the genus is considered undomesticated (Slavov et al. 2014). As a crop, there is a need to generate new broadly adapted genotypes suitable for a range of environments including both agricultural and marginal lands (Jing et al. 2012; Nijsen et al. 2012). This makes Miscanthus fit in well with the movement towards developing crops suited for marginal land so that fertile land is not taken away from food production (Cai et al. 2011; Donnelly et al. 2011). As such, the domestication and development of *Miscanthus* as a bioenergy feedstock can provide a case study of genetic resource utilization for novel crop development. However, for the full utilization of *Miscanthus* species in crop breeding an accurate taxonomy of *Miscanthus* is required.

*Miscanthus sensu lato* (s.l., in the broad sense) includes about 20 species depending on the author (Clayton and Renvoize 1986; Scally et al. 2001a,b; Clayton et al. 2006). However, recent molecular phylogenetics analyses have revised the genetic limits of the genus. Indeed, *Miscanthus sensu stricto* (in the strict sense) now includes only those species with a base chromosome number of 19 (Hodkinson et al. 2002b), with a resultant effect on the typification of *Miscanthus*.

Prior to the advent of molecular phylogenetics, the most comprehensive analysis of *Miscanthus* was that of Lee (1964a,b,c,d), who separated the genus into four sections: Kariyasua, Miscanthus, Triarrhena and Diandra. Section Diandra has now been excluded from Miscanthus due to sequence evidence (Hodkinson et al. 2002a), chromosome numbers and their possession of two anthers compared with three anthers in *Miscanthus s.s.* The African species formerly included in Miscanthus are better included in *Miscanthidium* (*M. ecklonii*, *M. junceus* and *M. sorghum* and *M. violaceum*) (Hodkinson et al. 2002a; Lloyd Evans et al. 2019) as they have a base chromosome number of 15. The South African species, *Miscanthus ecklonii* (syn *Miscanthus capensis*) was the type species for Miscanthus. However, as Hitchcock also defined *M. japonicus* (Trin.) Andersson (basionym *Saccharum floridulum* Labillardière; described in 1824), which is synonymous with *Miscanthus floridulus* (Labil.) as a second type for *Miscanthus*, then *M. floridulus* now becomes the type species for *Miscanthus s.s*.

Floral and physical characteristics can be variable, particularly as many species can hybridize. As a result, synonymy is very high in the genus. Indeed, the International Plant Names Index (IPNI, 2014) lists over 60 species, of which only 11–12 species can actually be recognized (https://www.kew.org/data/grasses-db/sppindex.htm#M).

For biofuel production, *Miscanthus* ×*giganteus* Greef et Deuter ex Hodkinson and Renvoize is the most interesting cultivar. This is posited as a hybrid of *M. sacchariflorus* and *M. sinensis* (Hodkinson and Renvoize 2001), which is triploid and sterile in a post-zygotic barrier that results from abnormal male and female gametophyte production (Słomka et al. 2012).

There has, however, been no large-scale analysis of the phylogenetic relationship of *Miscanthus* species, with the largest phylogram to date being an integrated image presented by Hodkinson et al. (2015) which was, itself based on three original studies Hodkinson et al. (2002a,b,c) and Swaminathan et al. (2010)

It should be noted that the exact origins of *Miscanthus* ×*giganteus* may be in doubt, as no complete chloroplast genomes have been published. Rather, *M.* ×*giganteus* has only been shown to have the plastid type of *M. sacchariflorus* (Hodkinson et al. (2002a,b) and De (2012).

*Miscanthus* species are C4 grasses, with high CO 2 use efficiency (Clifton-Brown and Jones 1997). Moreover, for a tropical plant Miscanthus shows a wide range of cold tolerance and certain *Miscanthus* ×*giganteus* cultivars can survive frost and winter freezing of below ground rhizomes (they are hardy to USDA zone 3) (Zuk et al. 2016).

*Miscanthus* is a member of the Saccharinae subtribe (as exemplified by sugarcane) and both species lie within the 3.4 million years of divergence where hybridization is possible (Lloyd Evans and Joshi 2016). Sugarcane and *Miscanthus floridulus* can hybridize in the wild and the introgression of *Miscanthus* into Saccharum (and *Saccharum* into Miscanthus) to produce miscanes focussing on biomass and high sucrose is a promising field of study to increase the biomass of *Miscanthus* and to introduce increased cold tolerance into sugarcane (Głowacka et al. 2016 and Burner et al. 2015). As such understanding the relationships of Miscanthus to Saccharum could have significant consequences for bioenergy crop development.

Using 13 long-range PCR primers previously developed for sugarcane chloroplast sequencing (Lloyd Evans et al. 2018), along with mining GenBank for assembled *Miscanthus* chloroplasts and GenBank’s Sequence Read Archive (SRA) for accessions where the chloroplastgenome could be assembled, a phylogeny consisting of 26 *Miscanthus*, two *Miscanthidium* and four *Saccharum* accessions was generated.

The recent finding that chloroplasts are completely expressed led to the attempt at assembling the chloroplast of *M. lutarioriparius* from transcriptomic data. This assembly is presented and integrated with the other whole *Miscanthus* chloroplasts assembled in this study.

The findings presented in this paper demonstrate that *Miscanthidium* is closer to *Saccharum* than *Miscanthus*. *Miscanthus* can be divided into three clades, represented by: *M. sacchariflorus* and *M. lutarioriparius*; *M sinensis*, *M. floridulus* and *M. transmorrisonensis*; and *M. oligostachyus*. *M. transmorrisonensis* and *M. floridulus* are mixed within the phylogeny, indicating that thy represent a single species. A single *Miscanthus sacchariflorus* accession lies as an outlier to *M. sinensis*, indicating that this may represent a previously unrecognized species. The divergence between *M. lutarioriparius* and *M. sacchariflorus* is sufficient for *M. lutarioriparius* (formally *M. sacchariflorus subsp lutarioriparius*) to be promoted to species level. The female ancestor of two *M.* ×*giganteus* accessions analysed in this study is shown to be *M. lutarioriparius* and not *M. sacchariflorus*.

## Materials and Methods

### Plant Samples and DNA Isolation

*Miscanthus sinensis* cv ‘Andante’, *Miscanthus transmorrisonensis* cv ‘Evening Maiden Grass’, *Miscanthus transmorrisonensis* Hayata, *Miscanthus sinensis* ‘Dixieland’, *Miscanthus sinensis* ‘Rigoletto’, *Miscanthus sinensis* var zebrinus, *Miscanthus sinensis* ‘Roland’ and *Miscanthus sacchariflorus* ‘Hercules’ were obtained from the Royal Botanic Gardens,Kew Seed Collection or Living Collection. *Miscanthus olygostachyus* var Afrika, *Miscanthus times giganteus* cv ‘Harvey’, *Miscanthus sinensis* var purpurascens and *Miscanthus sacchariflorus* ‘Robustus’ were obtained from a local nursery (Cambridge, UK). *Miscanthus floridulus* US56-022-03 and *Miscanthus sinensis* ssp. condensatus ‘Cabaret’ were obtained from the USDA collection. Seeds were grown to 8cm tall prior to harvest; otherwise young leaves were cut from established plants (5g total). DNA was extracted from seedlings and young leaves from whole plants using the standard CTAB method (Wang et al. 2010). Total DNA from *Miscanthus sinensis* ‘Andante’ was sequenced by Edinburgh Genomics using Illumina protocols and an Illumina HiSeq 4000 system. All the other samples had their chloroplast DNA amplified using 13 primers previously designed for *Saccharum* species amplification (Lloyd Evans et al. 2019). Amplicons were generated and sequenced by BeauSci Ltd, Cambridge, England using Illumina protocols and an Illumina HiSeq 4000 system.

### Whole Chloroplast Genome Assemblies

The *Miscanthus sinensis* cv ‘Andante’ chloroplast was assembled from whole genome reads, as described previously (Lloyd Evans and Joshi 2016) using the *Miscanthus floridulus* (LN869215.1) chloroplast assembly as a reference forbaiting and an untrusted reference for assembly. The plastome was assembled using Mirabait (Chevreux et al. 1999) and SPAdes v 3.10 (Bankevich et al. 2012), with a baiting k-mer of 27 and an assembly k-mer series of 25, 33, 55 and 77. The region corresponding to the second inverted repeat was copied, inverted and stitched into the genome (that this region represented a repeat was confirmed by increased coverage compared with the remainder of the genome).

Chloroplasts sequenced using PCR primers had raw reads adapter trimmed, prior to sequence cleaning with Trimmomatic (Bolger et al. 2014). Trimmed reads were assembled, typically generating seven or eight scaffolds, which could be arranged on the *Miscanthus sinensis* cv ‘Andante’ backbone. Any gaps in the initial assemblies were filled by excising a 1kb region around the gap prior to re-assembling this region by read baiting with Mirabait and assembling with SPAdes before incorporating the reassembled region back into the assembly.

The remaining chloroplast genomes were assembled from datasets submitted to the GenBank Sequence Read Archive (SRA). Downloaded datasets were as follows: *Miscanthus sinensis* ‘Gold Bar’ (SRR558867); *Miscanthus sinensis* ‘Nippon’ (SRR4029058); *Miscanthus sinensis* ‘Malepartus’ (SRR486616); *Miscanthus* × *giganteus* (SRR072880) and *Miscanthus sacchariflorus* cv ‘Hercules’ (SRR486748). In all cases, libraries over 10 Gb in size were reduced to 9 Gb by random selection using seqtk (https://github.com/lh3/seqtk/blob/master/README.md). All chloroplasts were assembled as described previously (Lloyd Evans and Joshi, 2016), using the newly assembled *Miscanthus sinensis* ‘Andante’ chloroplast as a reference.

The *Miscanthus lutarioriparius* chloroplast was assembled from transcriptomic data, using the SRA datasets: SRR5096715 and SRR2962583. The use of transcriptomic data for assembly of complete chloroplast sequences is a novel approach. Indeed, it was only recently that complete transcription of plant chloroplasts was reported (Shiet al. 2016). For the assembly of *Miscanthus lutarioriparius* transcriptomic data a variant of a previously published method was employed for our initial assembly, using the *Miscanthus sinensis* ‘Andante’ assembly as a reference. The genome assembled as 12 contigs, which could be joined together into three scaffolds. However, there were assembly issues with the rDNA cassette in the two inverted repeats. Transcriptomic data represents a combination of chloroplast transcripts, mitochondrial transcripts and genomic transcripts and these were all extracted from the read pool in our baiting step. However, chloroplast transcripts should occur at much higher frequency and chloroplast assemblies have ∼38.4% GC content (see supplementary Document 1). Using these criteria contigs derived from chloroplast transcripts could be extracted from the assembly pool and were stitched together manually to cover the entirety of the rDNA region in the chloroplast. This assembly was used as a template to re-construct this region using a k-mer of 31 and a minimum occurrence of 2 (n=2) in the baiting process. This time, the rDNA regions in the inverted repeats assembled completely, and were incorporated within the chloroplast.

### Assembly Finishing

To ensure that the assembly was of high quality, all assembled chloroplasts were finished and polished with a novel pipeline. Raw reads from the SRA pool were mapped back to the assembly with BWA (Li and Durbin 2009), tagging duplicate sequences with Picard tools (http://broadinstitute.github.io/picard), optimizing the read alignment with GATK (McKenna et al. 2010) and finally polishing and finishing with Pilon 1.2.0 (Walker et al. 2014).

### Miscanthus lutarioriparius Chloroplast Sequencing and Assembly

Subsequent to our RNA-based assembly of M. lutarioriparius, plants became available via Pan-Global Plants, Framption-on-Severn, England. Chloroplasts were isolated from total DNA using magnetic bead capture, as described previously (Lloyd Evans and Hughes 2020). Chloroplast DNA was sequenced with Oxford Nanopore Technologies MinION (Lloyd Evans and Hughes 2020), assembled and error corrected with Canu (Bankevich et al. 2012). Sequences for the other Miscanthus chloroplast genomes presented in this paper were also confirmed by MinION sequencing. The MinION data was also passed through the Organelle PBA pipeline (with minor modifications for MinION as opposed to PacBio data) (Soorni et al. 2017) and the relative proportions of chloroplasts with SSC regions in the forward and reverse orientations were determined. These data were also used to derive the two sequence variants of *Miscanthus* mitochondrial genomes.

### Chloroplast Annotation

The finished chloroplast alignments were orientated so that they finished with the IR_*B*_ region prior to batch upload to the Verdant annotation server (McKain et al. 2017) for automatedannotation. Annotation files were downloaded from Verdant in ‘.tbl’ (Verdant native) format. Genes and pseudogenes identified from a previous analysis (Lloyd Evans and Joshi, 2016), and which were not present in the Verdant annotation, were mapped with Exonerate (Slater and Birney 2005) and were placed in a separate ‘.tbl’ format file. A custom BioPerl script was written that integrated the Verdant output and the additional gene output along with data corresponding to gene annotation and cultivar level taxonomies prior to outputting an EMBL format file and a chromosome file that could be automatically uploaded to the EMBL database. All assembled chloroplasts were submitted to ENA under the project identifier PRJEB20532 and the ENA identifiers: ?????.

### GC Content Determination

As genomic ribosomal RNA reads can be a conflating influence in the assembly of chloroplasts from ribosomal RNA, reference GC contents were obtained from whole chloroplast genomes, whole mitochondrial genomes and complete nuclear genomes (see Supplementary Table 1). Fasta-formated sequences were downloaded or assembled (see supplementary Table 1 for sources) and were analyzed with the Cusp application of the EMBOSS package (Rice et al. 200).

### Whole Chloroplast Alignments

Whole plastid alignments were performed with SATÉ (version 2.2.2) (Liu et al. 2009), using MAFFT (Katoh and Standley 2013) as the aligner, MUSCLE (Edgar 2004) as the sub-alignment joiner and RAxML (Stamakis 2014) as the phylogenetic analysis application. Alignment iterations were run for 20 generations past the iterations that yielded the best likelihood score to ensure that the correct global alignment minimum had been reached. This optimal alignment was subsequently edited manually to remove any obvious errors and to trim all gaps of >20bp due to only a single sequence to 10bp (a list of all chloroplast assemblies with sequence and voucher accessions used in this study are given in Supplementary Table 1).

### Phylogenomic Analyses

The whole plastid alignment was divided into LSC, IR and SSC partitions. Bestfit evolutionary models for each partition were selected using PartitionFinder (Lanfear et al. 2014) and the AICc criterion. The LSC and SSC regions in the whole plastids best fit the GTR+Γ model and the IR regions best fit the GTR+Γ+I model. As the GTR+Γ and GTR+Γ+I models overlap, the proportion of invariant sites (‘I’) was omitted from all models (thisaccounts for the same phenomenon as the Gamma distribution and the simultaneous use of both parameters can cause convergence problems in Bayesian analyses). Bayesian analyses were run with MrBayes (version 3.1.2) (Ronquist and Huelsenbeck 2003) and Maximum Likelihood analyses were run with RAxML (version 8.1.17) (Stamakis 2006).

Bayesian Markov Chain Monte Carlo (MCMC) analyses were run with MrBayes 3.1.2, using four chains (3 heated and 1 cold) with default priors run for 15 000 000 generations with sampling every 100th tree. The whole plastid data were divided into LSC, IR_*B*_, SSC and IR_*A*_ partitions and model parameters were unlinked across partitions. For every partition the GTR+Γ model was employed. Two independent MrBayes analyses, each of two independent runs, were conducted. To avoid any potential over-partitioning of the data, the posterior distributions and associated parameter variables were monitored for each partition using Tracer v 1.6 (Rambaut et al. 2014). High variance and low effective sample sizes were used as signatures of over-sampling. Burn-in was determined by topological convergence and was judged to be sufficient when the average standard deviation of split frequencies was <0.001 along with the use of the Cumulative and Compare functions of AWTY (Nylander et al. 2008). The first 5 000 000 generations were discarded as burn-in, and the resultant tree samples were mapped onto the reference phylogram (as determined by maximum likelihood analysis) with the SumTrees 4.0.0 script of the Dendropy 4.0.2 package (Sukumaran and Holder 2010).

Maximum Likelihood inference and bootstrapping were performed in RAxML using the same partitioning scheme as detailed above, employing the GTR+Γ model for all partitions. To obtain the best tree RAxML was run without resampled replicates for 100 generations. The most likely tree was obtained in the 70th generation and this was used as the reference tree in all subsequent analyses. To confirm this topology, a second, independent run of RAxML with different seed parameters was also run. Both tree topologies were identical.

To provide support for the Maximum Likelihood phylogeny, a total of 12 500 bootstrap replicates were analysed. Replicate trees were summarized with SumTrees before being mapped onto the best maximally likely tree as determined above.

In addition to RAxML based analyses, the tree topology and SH-aLRT single branch supports were generated with IQ-Tree2 (Minh et al. 2020) employing 10 000 replicates.

### Divergence Time Estimates

Divergence times were estimated using BEAST 2.4.4 (Drummond et al. 2012), on a 16-core server running Fedora 25. The concatenated analysis (LSC, IR_*A*_, SSC, IR_*B*_) analysis was run for 50 million generations with sampling every 1000 replication under the GTR + Γ model with six gamma categories. The tree prior used the birth-death with incomplete sampling model (Drummond et al. 2012), with the starting tree being estimated using the unweighted pair group method with arithmetic mean (UPGMA). The site model followed an uncorrelated lognormal relaxed clock (Drummond et al. 2006). The analysis was rooted to *Zea mays* with the species’ age of origin given as a normal distribution describing an age of 13.8 ± 2 million years (Estep et al. 2014). Convergence statistics were estimated using Tracer v.1.5 (Rambaut et al. 2013) after a burn-in of 15 000 sampled generations. Chain convergence was estimated to have been met when the effective sample size was >200 for all statistics. Tree samples were integrated with SumTrees to generate the maximum clade credibility tree and to determine the 95% highest posterior density (HPD) for each node. The final tree was drawn using FigTree v.1.4.3 (http://tree.bio.ed.ac.uk/software/figtree/).

## Results

### Chloroplast Assembly

Of the 23 *Miscanthus* accessions analysed in this study, and despite employing both subset selection from whole genome data and assembly from PCR products there were no issues encountered with assembling the chloroplasts. Indeed, using the pipeline based on Mirabait, SPAdes, Verdant and Exonerate it typically took less than 2 hours to assemble and annotate a single chloroplast genome on a standard laptop (1 Tb hard drive, Intel core i5, 8 Gb RAM) running Linux. It also proved possible to completely assemble the *Miscanthus lutarioriparius* chloroplast from transcriptomic data. Assembled chloroplast genomes ranged in size from ???? bp to ???? bp, with the *Miscanthus sinensis* accessions ranging in size from ??? to ??? bp, the *Miscanthus sacchariflorus* accessions ranging in size from ??? to ???? bp. All *Miscanthus* accessions possessed 84 protein-coding genes, 33 non-protein coding genes (tRNAs and rRNAs), 3 pseudogenes and 4 origins of replication (discounting duplications). Exemplar assemblies, showing the range of *Miscanthus* species are shown in Figure 1.

**Figure 1:**
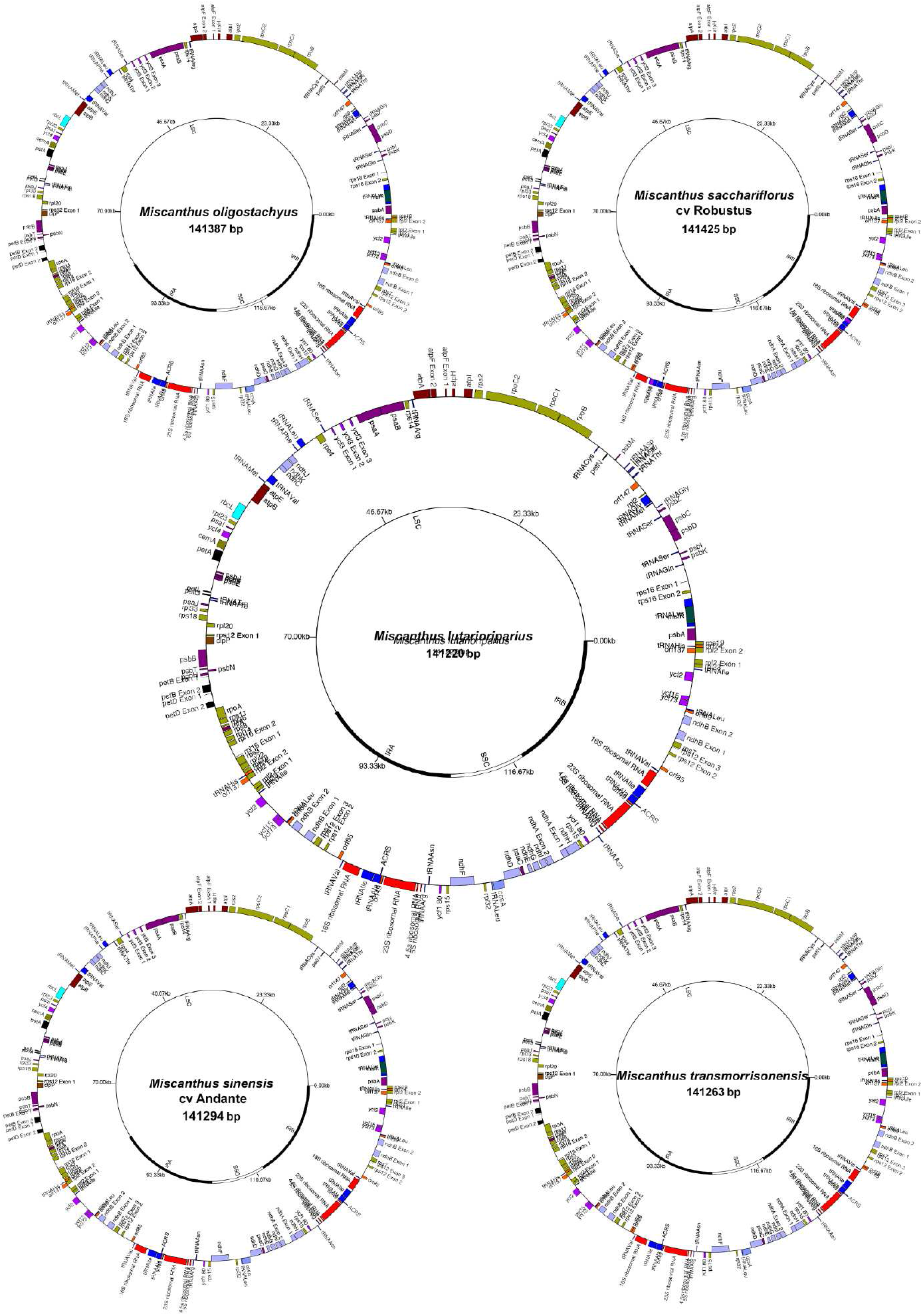
Schematic for the chloroplast assemblies of five reference *Miscanthus* speices. Schematic images showing the chloroplast assemblies and gene contents of five reference *Miscanthus* species: *Miscanthus oligostachyus, Miscanthus sacchariflorus, Miscanthus lutarioriparius, Miscanthus sinensis* and *Miscanthus transmorrisonensis.*

As now appears common in the majority of grasses, MinION Long read assembly revealed that the chloroplast genome in all the *Miscanthus* species analyzed existed as two distinct isoforms of equal sequence length, but with the short single copy (SSC) inverted between them (Soorni et al. 2017). Analysis (Table 1) revealed an almost equal distribution of the two sequence types, except for *Miscanthus oligostachyus* where the forward (canonical) SSC form predominated by almost 2:1. Both isoforms were annotated and submitted to the international sequence databases.

**Table 1:**
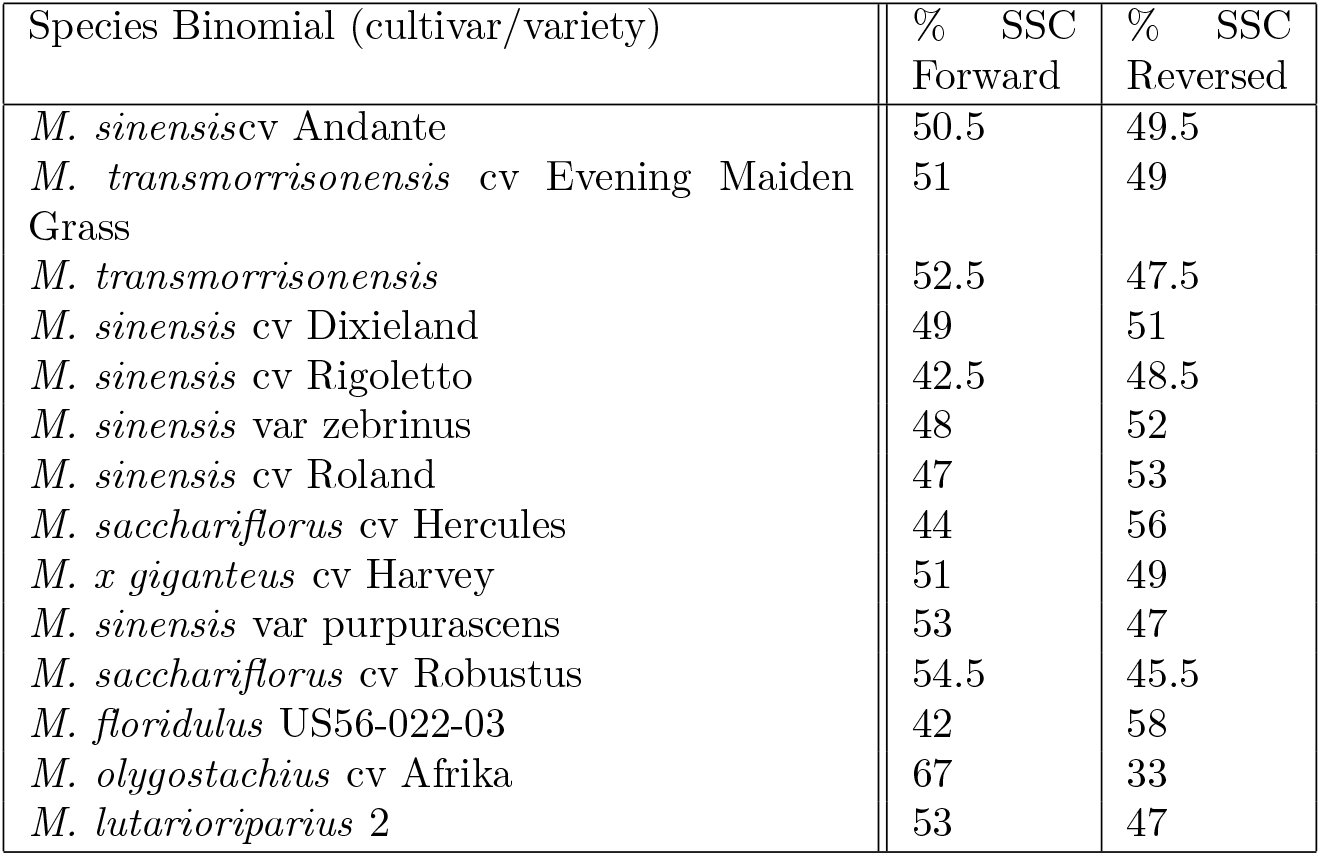
Table showing the Miscanthus species and cultivars analyzed along with the percentage of its chloroplast population with the Short Single Copy (SSC) region in the forward and reversed orientations.

| Species Binomial (cultivar/variety) | % SSC Forward | % SSC Reversed |
| --- | --- | --- |
| *M. sinensis*cv Andante | 50.5 | 49.5 |
| *M. transmorrisonensis* cv Evening Maiden Grass | 51 | 49 |
| *M. transmorrisonensis* | 52.5 | 47.5 |
| *M. sinensis* cv Dixieland | 49 | 51 |
| *M. sinensis* cv Rigoletto | 42.5 | 48.5 |
| *M. sinensis* var zebrinus | 48 | 52 |
| *M. sinensis* cv Roland | 47 | 53 |
| *M. sacchariflorus* cv Hercules | 44 | 56 |
| *M. x giganteus* cv Harvey | 51 | 49 |
| *M. sinensis* var purpurascens | 53 | 47 |
| *M. sacchariflorus* cv Robustus | 54.5 | 45.5 |
| *M. floridulus* US56-022-03 | 42 | 58 |
| *M. oligostachyus* cv Afrika | 67 | 33 |
| *M. lutarioriparius* 2 | 53 | 47 |

### Phylogenomic Analyses

Both RAxML and IQ-Tree yielded the same tree topology, indicating general robustness. The backbone of the tree is well supported, with some terminal nodes receiving much less support (to be expected as many of the sequences were nearly identical).

Phylogenetic analyses reveal that *Miscanthus s.s*. is monophyletic and is sister to a clade formed by Saccharum and Miscanthidium (with 100% support). The phylogeny presented here strongly supports the separation of *Miscanthidium* from *Miscanthus*, with *Miscanthidium* being sister to *Saccharum* rather than sister to *Miscanthus* (Figure 2). Within *Miscanthus* there is an early divergence of a clade formed from *Miscanthus lutarioriparius* and *Miscanthus sacchariflorus* from the remaining *Miscanthus* species. *Miscanthus olygostachius* forms an outgroup to a clade formed from *Miscanthus sinensis, Miscanthus transmorrisonensis* and *Miscanthus fioridulus.* However, *Miscanthus floridulus* and *Miscanthus transmorrisonensis* accessions are mixed within the phylogeny.

For clarity, within Figure 2 the relationships between the crown species and *Miscanthus sacchariflorus* are shown inset. This indicates that *Miscanthus sinensis* is divided into two distinct subspecies, one that includes *Miscanthus sinensis* Malepartus and the other which includes *Miscanthus sinensis* Condensatus cv Cabaret. Thus is appears that Malepartus and Condensatus refer to two distinct subspecies of *Miscanthus sinensis*.

**Figure 2:**
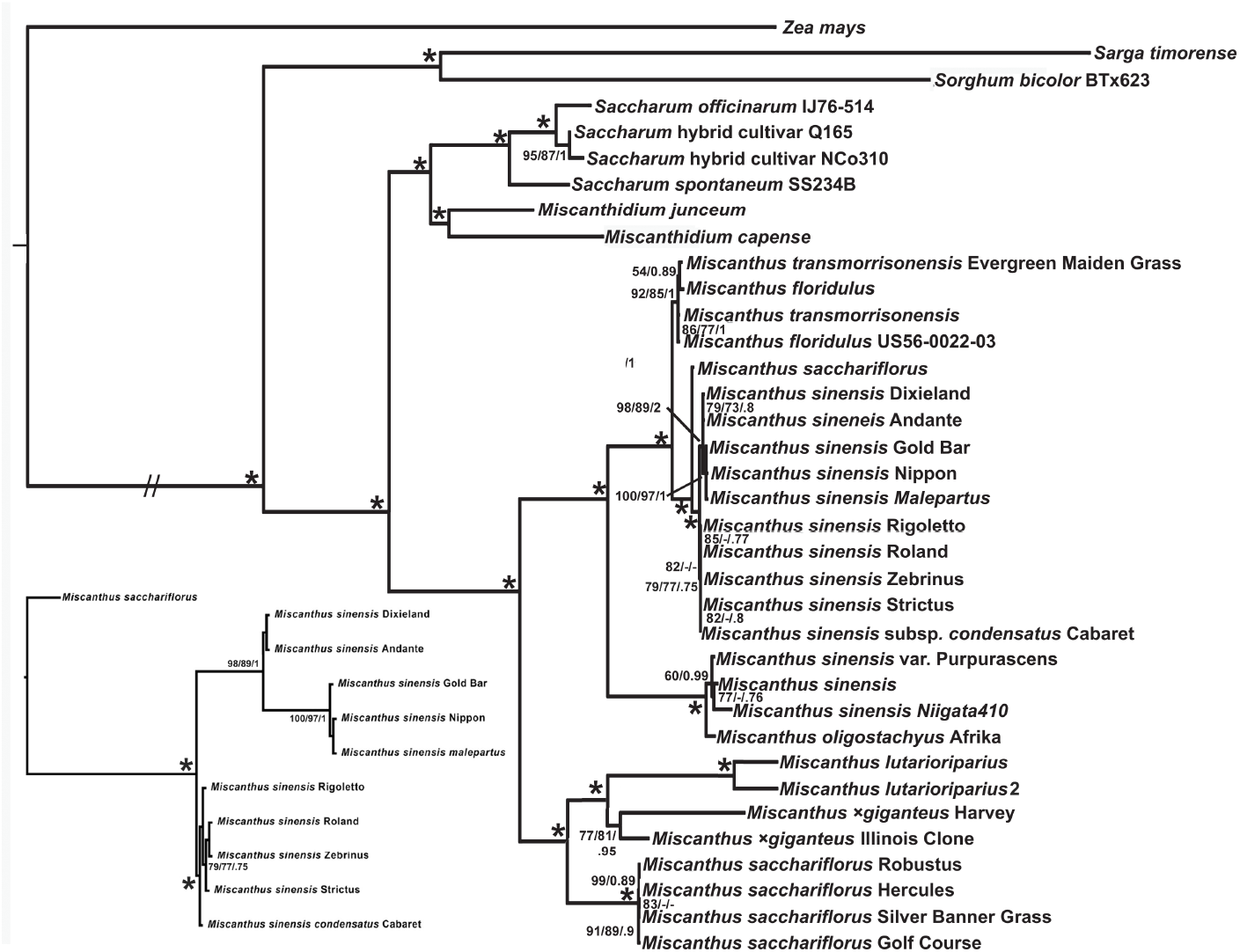
Phylogram showing the evolution of all the main recognized *Miscanthus* speices. Phylogram comparing the evolution of the main *Miscanthus* species and types. *Zea mays* is employed as an outgroup and the sister relationship of *Miscanthus* to *Miscanthidium* and *Saccharum* is shown. The relationship between *Miscanthus sinensis* malepartus and *Miscanthus sinensis* condensatus is shown in more detai, inset. Numbers at branches reveal non-parametric bootstrap values (>70%) and Bayesian inference values (≥0.75). // represents a long branch that has been shrunk by 66% to better reveal the relationships at the crown of the tree. Asterisks (*) represent nodes with 100% support, whereas a dash (−) represent branch support that is lower than the cutoff. Three branch supports, SH-aLRT single branch tests/non-parametric bootstrap and Bayesian Inference are shown.

The DNA and RNA based *M. lutarioriparius* assembles are sisters in Figure 2 and both emerge as sister to *M.* × *giganteus*. This shows the utility of assembling whole chloroplasts from RNA datasets in phylogenetic analysis.

A separate and distinct outgroup (with 100% support) to *Miscanthus floridulus* + *Miscanthus transmorrisonensis* and the core *Miscanthus sinensis* accessions by *Miscanthus sinensis* var Purpurascens, a *Miscanthus sinensis* accession from Korea, *Miscanthus sinensis* cv Niigata410 and *Miscanthus oligostachyus* cv Africa.

Two accessions of *Miscanthus lutarioriparius* (from RNA assembly and DNA sequencing) are sister to *Miscanthus* × *giganteus*. This grouping is sister to *Miscanthus sacchariflorus* and forms an ougtgroup to all the remaining *Miscanthus* accessions.

### Timing of Evolutionary Events

The chronogram, Figure 3, which is calibrated to the evolutionary origins of *Zea mays*. This places the split between *Zea* and *Sorghum* at about 6.77 million years ago, which is consistent with our previous studies (Lloyd Evans and Joshi 2016). With more species within *Miscanthus* in this study, the last common ancestor of *Miscanthus* and *Saccharum* was 4.49 million years ago, almost 1.5 million years more ancient than our previous estimate (Lloyd Evans and Joshi 2016). The *M. lutarioriparius*/*M. sacchariflorus* grouping diverged from the remaining *Miscanthus* species 2.78 million years ago. Our novel *M. olygostachius* grouping last shared a common ancestor with *M. floridulus* and *M. sinensis* 1.46 million years ago, which is the same time that *Saccharum sinense* last shared a common ancestor with] *S. officinarum*.

**Figure 3:**
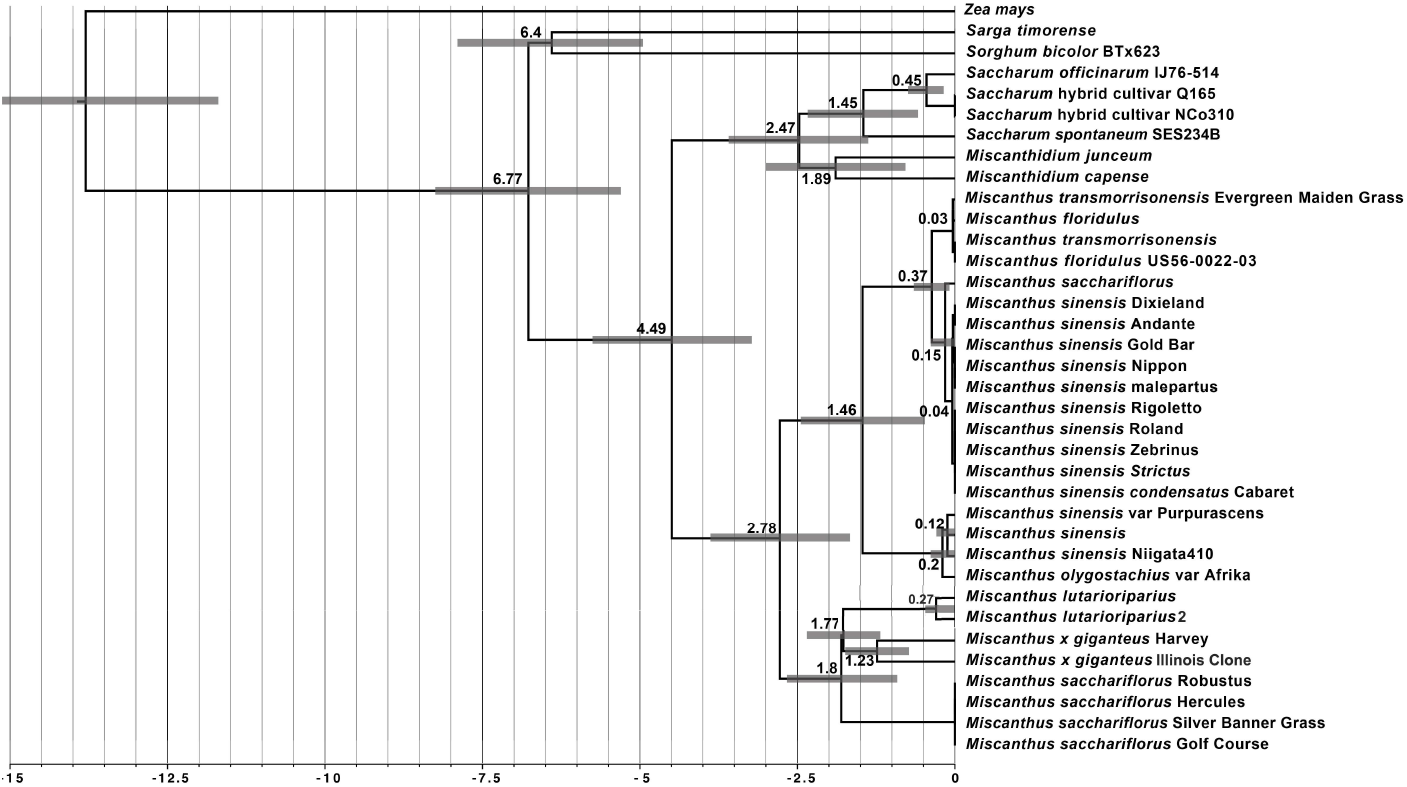
Chronogram derived from Beast analysis of the alignment yielding the phylogeny in Figure 2. Vertical bars and numbers at the base represent millions of years before present. Numbers at nodes represent the age of the node in millions of years before present and bars at nodes represent the 95% highest probability distribution of node ages.

The grouping of *M. floridulus* and *M. transmorrisonensis* last shared a common ancestor 370 000 years ago. The two subgroups of *M. sinensis* diverged 150 000 years ago.

## Discussion

*Miscanthus* has the potential to be a very valuable biofuel crop, particularly as many species are cold tolerant. This allows *Miscanthus* to grow outside the tropical zone, making it a potentially more widespread crop than sugarcane (Saccharum hybrids). In addition, *Miscanthus* is generally less dependent on water supply and is more nutrient (particularly nitrogen) efficient than sugarcane (Anderson et al. 2014), which mans that it can, potentially, be grown on marginal land (Kosinkova 2015). With electricity stations that can be powered solely using biofuel generators that are now coming online (Gough 2018) this makes *Miscanthus* an attractive bioenergy/biofuel crop in warm-temperate and temperate regions of the world.

However, Miscanthus has not yet been fully domesticated and there has been no large-scale phylogeny of Miscanthus species based on a large number of characters. The current analysis, using complete chloroplasts (with an alignment length of ????? bp) and ???? accessions is the largest and most comprehensive phylogenomic study of Miscanthus species performed to date.

The phylogeny (Fig. 2) and chronogram (Fig. 3) reveal that *Miscanthidium* species are clearly distinct from *Miscanthus* species and are actually closer to *Saccharum* species (being ??? million years divergent). *Miscanthus* as a genus is monophyletic (100% support) and is divided into three main clades.

In contrast to the combined phylogeny presented by Hodkinson et al. (2015) the chloroplast phylogeny places the clade formed by *Miscanthus sacchariflorus* and *Miscanthus lutarioriparius* as the earliest diverging (the Hodkinson et al. (2015) paper places a *Miscanthus olygostachyus* clade as the earliest diverging). The phylogeny presented in this paper also shows clear evolutionary divergence between *Miscanthus sacchariflorus* and *Miscanthus sacchariflorus* subsp. lutarioriparius (L. Liu ex S.L. Chen & Renvoize) Q. Sun & Q. Lin. The 1.8 million year divergence between *M. sacchariflorus* and *M. sacchariflorus* subsp. lutarioriparius supports the elevation of *M. lutarioriparius* to species level (as described for *Miscanthus lutarioriparius* L. Liu ex S.L. Chen & Renvoize). Moreover, the phylogeny presented in this paper places *M. lutarioriparius* rather than *M. sacchariflorus* as the female parent for *M. times giganteus*.

The next clade is bounded by *Miscanthus olygostachyus* at its root. Interestingly, this species is sister to *Miscanthus sinensis* var Purpurascens and two previously published sequences of *M. sinensis* and *Miscanthus floridulus* (Lloyd Evans and Joshi 2016; Nah et al. 2016; Tsuruta et al. 2017). This would indicate that the *M. sinensis* forms have either been misidentified or they represent hybrids of *M. oligostachyus* with *M. sinensis*. Due to the nature of chloroplast phylogenetics, we demonstrate here that *M. oligostachyus* is the female parent. This potentially adds *Miscanthus* × *oligostachyus* to the list of *Miscanthus* hybrids, though more work at the 45s ribosomal RNA and/or low copy number gene level will be needed to confirm that *Miscanthus* sinensis var purpurascens, *M. sinensis* (GenBank: NC 028721.1) and the other *M. sinensis* accessions are truly hybrids.

Crown *Miscanthus* species are formed from two sister clades. The first clade is formed from *M. transmorrisonensis* and *M. floridulus* accessions. As both species are intermixed in the phylogeny this strongly indicates that they are a single species, diverging from *M. sinensis* some 370 thousand years ago. *Miscanthus floridulus* (Labill.) Warb. ex K. Schum. & Lauterb. was formally published in 1901 (Lauterback and Schumann, 1901). *M. transmorrisonensis* Hayata was formally published in 1911 (Hayata, 1911). This means that, excluding hybridity, both species can be collapsed into *M. floridulus*. Thus *Miscanthus* as a genus retains its type species of *M. floridulus*.

A previously published *M. sacchariflorus* chloroplast assembly (GenBank: NC 028720.1) serves as an outgroup for the *M. sinensis* clade. This accession, however lies as sister to the clade formed by *M. sacchariflorus* + *M. sinensis*, as such, it cannot be a true *M. sacchariflorus* and could, potentially, represent a novel species. However, the other *M. sinensis* accessions are all very closely related,indicating the small genetic variation between *M. sinensis* accessions held in Europe and America. *M. sinensis* × *M. sacchariflorus* hybrids are not unknown, however (Tamura et al. 2016) and this accession could represent a hybrid with an early diverging *M. sinensis* accession.

Within the core *M. sinensis* clade, *M. sinensis* ‘Dixieland’ *M. sinensis* ‘Andante’, *M. sinensis* ‘Gold Bar’ and *M. sinensis* ‘Nippon’ along with *M. sinensis* malepartus form a discreet clade with good support at the root. All these cultivars therefore probably belong to the *Miscanthus sinensis* subspecies malepartus, whilst the remaining two accessions (Andante and Dixieland) belong to the subspecies *Miscanthus sinensis* subspecies sinensis. However, the phylogeny presented in Figure 2 demonstrates that the *M. sinensis* subsp. condensatus accessions studied are essentially clonal (with little or no variation in their chloroplast sequences) and little branch support in the phylogeny beyond the basal node. It should also be noted that the remaining Miscanthus sinensis accessions (cultivars Rigoletto, Roland, Zebrinus, Strictus and var condensatus cv Cabaret) all form a monophyletic clade which is 100% supported at the root, but which has poor internal branch support, indicating a high degree of chloroplast similarity. The root of this clade is *Miscanthus sinensis* var condensatus cv Cabaret. This clade is distinct from both the *Miscanthus sinensis* subsp sinensis and *Miscanthus sinenis* subsp malepartus clades, giving strong support we have a separate *Miscanthus sinensis* condensatus subspecies. The current accepted name is *Miscanthus sinensis* var. condensatus (Hack.) Makino, but the data presented in this article demonstrates that this is a true subspecies and should more correctly be referred to as *Miscanthus sinensis* subsp. *condensatus* (Hack.) T. Koyama.

## Conclusion

The largest whole chloroplast study on genus *Miscanthus* ever performed is presented. This study clearly demonstrates the distinction between *Miscanthus* and *Miscanthidium*, with the genera diverging 4.49 million years ago. Indeed, from chloroplast data and also based on our recent 45s ribosomal RNA analysis (Lloyd Evans and Joshi 2020) *Miscanthidium* is closer to *Saccharum* than to *Miscanthus*. As well as assembling 22 *Miscanthus* accessions from DNA short read data, the whole chloroplast of *M. lutarioriparius* is assembled from transcriptomic data, providing strong supporting evidence that the chloroplast of grasses are transcribed in their entirety.

A clade formed from *M. lutarioriparius*, *M. sacchariflorus* and *M.* × *giganteus* is the earliest diverging of the Miscanthuses and is 2.78 million years divergent from a clade formed from *M. olygostachius*, *M. sinensis*, *M. transmorrisonensis* and *M. floridulus*. *M. lutarioriparius* is sister to *M. sacchariflurus*, with the species diverging 1.8 million years ago. Interestingly, the chloroplast phylogeny places *M. lutarioriparius* rather than *M. sacchariflorus* as the female parent of two *M.* × *giganteus* accessions. These data demonstrate that *M. lutarioriparius* should be elevated to species level.

*Miscanthus* × *giganteus* is known from Japan, but *Miscanthus lutarioriparius* is endemic to the middle and low altitudes in the Yangzi River area of China (Li et al. 2013). *Miscanthus sinensis* and *Miscanthus sacchariflorus* are both native to Japan, which is why it was always believed that *M.* × *giganteus* was a hybrid of *M. sinensis* and *M. sacchariflorus*. However, the phylogenetics presented in this paper indicate that *M.* × *giganteus* has a *Miscanthus lutarioriprius* type chloroplast and may actually be a hybrid of *M. lutarioriparius* and *M. sinensis*. However, the *M. sinensis* and *M. lutarioriparius* chloroplast types last shared a common ancestor 1.77 million years ago. This either means that the original hybrid leading to *M* × *giganteus* is truly ancient or that there is a cryptic species involved that is closer to *M. lutarioriparius* than it is to *M. sacchariflorus*.

*M. olygostachius* is sister to and 1.46 million years divergent from a clade formed by *M. sinensis*, *M. floridulus* and *M. transmorrisonensis*. *M. olygostachius* is sister to three accessions previously identified as *M. sinensis*, indicating that these may represent hybrids.

Within the crown Miscanthus clade, a group formed by *M. transmorrisonensis* and *M. floridulus* is sister to the remaining *M. sinensis* accssions. *M. transmorrisonensis* and *M. floridulus* are not separated within the phylogeny, indicating that they represent a single species, with *M. floridulus* gaining taxonomic precedence as the name of the group.

An accession named *M. sacchariflorus* is the outgroup to all the *M. sinensis* accessions, indicating that this may have been miss-identified. The *M. sinensis* group is split into three clades (all the clades having good support at the root), representing subspecies sinensis, subspecies malepartus and subspecies condensatus within the *Miscanthus sinensis* clade. This indicates that cultivars Rigoletto, Roland, Strictus and Zebrinus should classified as *Miscanthus sinensis* subsp. condensatus.

## 1 Availability of data and materials

All sequences generated in this study have been submitted to ENA under the project accession ???. Sequence alignments and base phylogenies are available from Zenodo (DOI). Computer code developed for the project is available from GitHub: https://github.com/gwydion1/bifo-scripts.git.

## 4 Authors’ contributions

DLlE secured the funding, conceived the experiment, performed the analysis and historical research, analyzed the data and wrote the paper.

## 5 Acknowledgements

I would like to thank Mr B. Hughes for performing the MinION sequencing.

