## Supplementary Document 1 for "Assembly of 23 Plastid Genomes Provides the Chloroplast View on *Miscanthus* Origins"

Dyfed Lloyd Evans

June 28, 2020

### GC Content Analysis in Chloroplast, Mitochondria and Genomes

| Species | Voucher or Accession | GenBank Accession | Reference | % GC Content |
|---|---|---|---|---|
| *Sorghum bicolor* | BTx623 | EF115542.1 | Saski et al. 2007 | 38.5 |
| *Sarga timorense* | | KF998272.1 | Kepers et al. (submitter) | 38.5 |
| *Zea mays* | B73 | AY928077.1 | Schnable et al. 2009 | 38.5 |
| *Saccharum officinarum* | IJ76-514 | LN849913 | Lloyd Evans & Joshi 2016 | 38.5 |
| *Saccharum* hybrid cultivar | Q165 | LN896359.1 | Lloyd Evans & Joshi 2016 | 38.4 |
| *Saccharum* hybrid cultivar | SP80-3280 | AE009947.2 | Calsa Jr et al. 2004 | 38.4 |
| *Saccharum spontaneum* | SES234B | LN849912.1 | Lloyd Evans, D. (submitter) | 38.4 |
| *Miscanthidium junceum* | | LN869216 | Lloyd Evans, D. (submitter) | 38.4 |
| *Miscanthidium capense* | | PRJEB17861 | Lloyd Evans, D. (submitter) | 38.4 |
| *Miscanthus floridulus* | PI295762 | LN869215.1 | Lloyd Evans and Joshi 2016 | 38.4 |
| *Miscanthus sinensis* | Andante | LM735682 | Lloyd Evans, D. (submitter) | 38.4 |
| *Miscanthus sacchariflorus* | Hercules | LN869218.1 | Lloyd Evans, D. (submitter) | 38.4 |

Table 1: Analysis for GC content in whole chloroplast sequences for a range of *Miscanthus* and related genera.

| Species | Voucher or Accession | GenBank Accession | Reference | % GC Content |
|---|---|---|---|---|
| *Saccharum officinarum* | Khon Kaen 3 | LC107874.1 LC107875.1 | Shearman et al. 2016 | 43.82 |
| *Sorhum bicolor* | BTx623 | DQ984518.1 | Allen et al. 2007 | 43.73 |
| *Zea mays* | B73 | AY928077.1 | Schnable et al. 2009 | 38.5 |
| *Tripsacum dactyloides* | Pete | NC_008362.1 | Allen et al. 2007 | 43.93 |
| *Zea mays* | B73 | NC_007982.1 | Clifton et al. 2004 | 43.93 |
| *Zea perennis* | | DQ645538.1 | Allen et al. 2007 | 38.4 |
| *Miscanthus sinensis* | Andante | [Partial Assembly] | Lloyd Evans et al. 2019 | 43.85 |

Table 2: Analysis for GC content in whole mitochondrial genomes for a range of *Miscanthus* and related genera. Note that the M. sinensis assembly is partial, but still of good enough quality to yield a % GC estimation.

| Species | Voucher or Accession | GenBank Accession | Reference | % GC Content |
|---|---|---|---|---|
| *Sorhum bicolor* | BTx623 | EnsEMBL | Paterson et al. 2008 | 41.4 |
| *Zea mays* | B73 | EnsEMBL | Schnable et al. 2009 | 47.2 |
| *Saccharum hybrid* | SP80-3280 | GenBank | Lloyd Evans et al. 2019 | 42.7 |
| *Miscanthus sinensis* | IGR-2011-001 | Phytozome | Miscanthus sinensis v7.1 DOE-JGI | 43.6 |

Table 3: Analysis for GC content in whole nuclear genomes for a range of *Miscanthus* and related genera. The data for sugarcane were derived from a combinationi of partial assemblies and contigs.
