## Supplementary Table 1 for "Assembly of 23 Plastid Genomes Provides the Chloroplast View on *Miscanthus* Origins"

Dyfed Lloyd Evans

June 28, 2020

### Sources of External Chloroplasts used in Phylogenetics

| Species | Voucher or Accession | GenBank Accession | Reference |
|---|---|---|---|
| *Zea mays* | B73 | AY928077.1 | Schnable et al. 2009 |
| *Sorghum bicolor* | BTx623 | EF115542.1 | Saski et al. 2007 |
| *Sarga timorense* | | KF998272.1 | Kepers et al. (submitter) |
| *Saccharum officinarum* | IJ76-514 | LN849913 | Lloyd Evans & Joshi 2016 |
| *Saccharum* hybrid cultivar | Q165 | LN896359.1 | Lloyd Evans & Joshi 2016 |
| *Saccharum* hybrid cultivar | Nco310 | AP006714 | Asano et al. 2004 |
| *Saccharum spontaneum* | SES234B | LN849912.1 | Lloyd Evans, D. (submitter) |
| *Miscanthidium junceum* | | LN869216 | Lloyd Evans, D. (submitter) |
| *Miscanthidium capense* | | PRJEB17861 | Lloyd Evans, D. (submitter) |
| *Miscanthus floridulus* | PI295762 | LN869215.1 | Lloyd Evans and Joshi 2016 |
| *Miscanthus sinensis* | Andante | LM735682 | Lloyd Evans, D. (submitter) |
| *Miscanthus sacchariflorus* | Hercules | LN869218.1 | Lloyd Evans, D. (submitter) |
| *Miscanthus sinensis* | Niigata 410 | LC160131 | Tsuruta et al. 2017 |
| *Miscanthus sinensis* | | KR822688 | Nah et al. 2016 |
| *Miscanthus sacchariflorus* | | KR833687 | Nah et al. 2016 |

Table 1: List of all externally assembled chloroplasts employed in the phylogenomic analyses.
